# A 16-colour spectral flow cytometry panel to characterise T cell immunophenotypes in canine oral melanoma

**DOI:** 10.64898/2026.08.25.746237

**Authors:** E Hindriks, E l Lozano-Andrés, A Roos, MMJM Zandvliet, EJAM Sijts, F Broere

## Abstract

Advances in immunophenotyping of tumour-infiltrating lymphocytes (TILs) have improved our understanding of prognostic biomarkers and immune targets in human melanoma. However, whether the tumour-immune landscape in canine oral malignant melanoma (COMM) is concordant with that of human melanoma has not been properly defined. To address this gap, we developed a 16-colour spectral flow cytometry panel to characterise TIL phenotypic and functional profiles in COMM. Validation using mitogen-stimulated peripheral blood mononuclear cells from healthy dogs (*n* = 5) demonstrated robust identification of major T cell lineages, including regulatory T cells (T_regs_) and memory subsets, and reliable evaluation of their activation and exhaustion status. COMM patients (*n* = 8) displayed a distinct protumour microenvironment, characterised by an increased proportion of T_regs_, enrichment of tumour-specific exhausted-like T cells co-expressing programmed cell death protein 1 (PD-1) and tumour necrosis factor receptor 2 (TNFR2), and a reduction in cytotoxic CD8_⁺_ and natural killer T (NKT) cell populations compared with tissue-resident (*n =* 4) and circulating (*n* = 8) lymphocytes. Analysis of additional solid tumours, including a mast cell tumour, nerve sheath tumour and adrenal cortical carcinoma, further supported the capability of the panel to identify similar patterns of immune dysregulation across diverse canine tumour landscapes. Collectively, this work describes the first detailed evaluation of canine TIL immunophenotyping using spectral flow cytometry and provides insights into the immunosuppressive mechanisms shaping the tumour microenvironment in COMM. These findings not only increase our understanding of canine tumour immunology but also identify potential immune targets and support ongoing comparative immuno-oncology efforts.

## 1 Introduction

Emerging immunotherapies have significantly changed the treatment of various cancers, most notably metastatic melanoma (Regan et al., 2016). These therapies include immune checkpoint inhibitors (ICIs), which block inhibitory signals on immune cells, particularly tumour-infiltrating lymphocytes (TILs), to enhance the immune response against tumour cells (Rui et al., 2023). However, response rates in tumour treatment are variable and immune-related adverse effects are common; unfortunately, both are currently hard to predict (Lin et al., 2016). Therefore, a deeper understanding of the phenotype and functionality of TILs is needed.

Canine oral malignant melanoma (COMM) is a highly aggressive tumour with strong parallels to human mucosal melanoma in terms of histopathology, metastatic behaviour and immune evasion (Palma et al., 2021). This spontaneously occurring tumour in an immunocompetent animal makes COMM a comparative model to evaluate novel immunotherapies, benefiting both human and canine patients. However, the tumour micro-environment (TME) has been poorly characterised in dogs compared to that in humans.

In humans, T cells have been demonstrated to play a crucial role in antitumour immunity; their effector, memory, regulatory, and exhaustion status provide insight into tumour progression, immune evasion and therapeutic response (Kraja et al., 2025). Effector CD8^+^ T cells are directly involved in tumour cell killing after antigen recognition through MHC class I, whereas effector CD4^+^ T cells are indirectly involved in antitumour immunity by coordinating immune cell functions after antigen recognition through MHC class II (Lopez de Rodas et al., 2025). Memory T cells, particularly in tissues, are essential for a durable antitumour response and can be identified by markers such as CD45RA, CD62L, CCR7, and CD103 (Gavil et al., 2024). In contrast, regulatory T cells (T_reg_) suppress the antitumour response and are characterised by the expression of CD4, CD25, and FoxP3 (Tanaka and Sakaguchi, 2017). Exhausted T cells (T_ex_) are also immunosuppressive; they upregulate inhibitory molecules, whose signalling represses effector functions (Zebley et al., 2024). The discovery of T_ex_ has led to the development of ICIs, aiming to reinvigorate them by blocking their inhibitory molecules, such as programmed cell death (PD-1), its ligand (PD-L1), or cytotoxic T lymphocyte-associated protein 4 (CTLA-4).

In dogs, much less is known about TIL heterogeneity, including their activation, memory, regulatory, and exhaustion status. Current studies have focused on identifying CD4^+^ and CD8^+^ T cells and T_reg_ in canine melanoma (Sparger et al., 2021; Tominaga et al., 2010; Yasumaru et al., 2021) and mammary carcinoma (Estrela-Lima et al., 2010). Assessing TIL phenotypes is challenging due to the limited available methods. Immunohistochemistry (IHC) provides valuable spatial information (e.g. intratumoral versus stromal TILs) but is generally restricted to a single marker per tissue section (Muscatello et al., 2022). In contrast, flow cytometry enables high-dimensional single-cell analysis for extensive TIL immunophenotyping, though spatial information is lost. Conventional flow cytometry, however, is constrained in multiplexing by the number of available lasers, dedicated detectors and spectral overlap (Nolan, 2022). Spectral flow cytometry addresses this limitation by capturing the full emission spectrum of each fluorochrome and using mathematical unmixing to resolve overlapping spectral signatures. More similar fluorophores can be included in a single panel, thereby supporting larger panels and overcoming the limited availability of fluorophore-conjugated species-specific antibodies. Consequently, spectral flow cytometry enables more in-depth immunophenotyping and is currently being adopted in the veterinary field (McDonald et al., 2025).

In this study, we describe the development and validation of a 16-colour spectral flow cytometry panel specifically designed to characterise T cell immunophenotypes in COMM and other solid tumours. The panel includes lineage, activation, memory and exhaustion markers. Our objectives in this study are to develop and validate a robust spectral flow cytometry panel suited for canine immuno-oncology, map the distribution of different T cell phenotypic and functional profiles, and compare tumour-infiltrating, tissue-resident and circulating T cell phenotypes. By enabling high-dimensional single-cell analysis of canine TILs, our panel offers the possibility to investigate the effects of veterinary immunotherapies targeting the TME, providing translational insights for comparative immuno-oncology.

## 2 Materials and methods

### 2.1 Tissue digestion

Tumour biopsies were collected with owner consent at the University Clinic for Companion Animals, Faculty of Veterinary Medicine, Utrecht University; Evidensia Animal Hospital in Nieuwegein; or Veterinary Referral Centre in Gouda. Healthy oral mucosa biopsies were collected from four different dogs that were sacrificed at the end of an unrelated study approved by the Utrecht University committee for Experiments on Animals. Tissues were cut into small pieces (<2 mm) and digested for 30 min at 37 °C in Roswell Park Memorial Institute (RPMI)-1640 GlutaMAX (Gibco) containing 1 mg/ml collagenase IV (Sigma-Aldrich). Single cells were passed through a 70 μm nylon mesh strainer, washed with cold RPMI-1640 GlutaMAX culture medium supplemented with 5% foetal bovine serum (FBS; Bodinco) and 50 U/ml penicillin and 50 µg/ml streptomycin (Gibco), hereafter referred to as RPMI/5% FBS, and centrifuged for 10 min at 300 × g. Red blood cells were lysed with sterile Ammonium-Chloride-Potassium (ACK) lysis buffer (pH 7.4), consisting of Milli-Q water with 0.15 M NH_4_Cl, 10 mM KHCO_3_, and 0.1 mM ethylenediaminetetraacetic acid (EDTA; Invitrogen), for 2 min on ice. Single cells were resuspended in RPMI/5% FBS and freeze medium, consisting of 80% FBS and 20% dimethyl sulfoxide (DMSO; Honeywell), in a 1:1 ratio and cryopreserved at −130 °C until use.

### 2.2 Peripheral blood mononuclear cell isolation

Buffy coats, as residual material from clinical blood donations for transfusion purposes, were obtained with owner consent from healthy client-owned dogs at the University Clinic for Companion Animals, Faculty of Veterinary Medicine, Utrecht University. Peripheral blood mononuclear cells (PBMC) were isolated using density gradient centrifugation at room temperature (RT). Briefly, samples were diluted 1:1 with Dulbecco’s phosphate-buffered saline (DPBS; Corning), layered onto Ficoll-Paque PLUS (Cytiva), and centrifuged for 30 min at 800 × g with low acceleration and break. The cloudy interphase was transferred to a new tube and washed in cold RPMI/5% FBS, and centrifuged for 10 min at 450 × g with medium acceleration and brake. Cells were washed twice and centrifuged for 10 min at 250 × g with maximum acceleration and brake. Similar to tumour cells, PBMC were cryopreserved at −130 °C until use.

### 2.3 T cell proliferation assay

Cryopreserved PBMC from five different healthy donors were thawed in a water bath and washed with RPMI/5% FBS and centrifuged for 5 min at 300 × g. Next, cells were washed and resuspended in serum-free X-VIVO (Lonza Bioscience) culture medium supplemented with 0.05 mM β-mercaptoethanol (Sigma-Aldrich) and 50 U/ml penicillin and 50 µg/ml streptomycin. Cells were seeded at 2 × 10^5^ cells/well in a 96-well U-bottom plate in the presence or absence of 1 μg/ml phytohemagglutinin (PHA; Calbiochem) and incubated for 72 h at 37 °C and 5% CO_2_.

### 2.4 Flow cytometry staining

The antibodies and fluorophores used are listed in Table 1. Freshly thawed or cultured samples were washed with DPBS, centrifuged for 5 min at 300 × g and incubated with ViaKrome808 in DPBS for 30 min at 4 °C. Cells were washed with FACS buffer, consisting of DPBS supplemented with 2% FBS and 2 mM EDTA, and blocked with 5% healthy dog and goat serum in FACS buffer for 15 min at 4 °C. Next, the unconjugated antibodies against CD45, CD45RA and tumour necrosis factor receptor 2 (TNFR2) were added for 30 min at 4 °C. After washing, cells were incubated with the secondary antibodies in FACS buffer for 30 min at 4 °C. Cells were blocked with 5% normal mouse and rat serum in FACS buffer for 15 min at 4 °C, washed, and incubated with directly conjugated antibodies against PD-1 and CD62L in FACS buffer for 30 min at RT. Cells were rewashed and incubated with directly conjugated antibodies against CD21, CD4, CD8, CD94, and CD25 in FACS buffer for 30 min at 4 °C. Next, cells were fixed and permeabilised using the FoxP3 / Transcription Factor Staining Buffer Set (eBioscience) according to the manufacturer’s protocol and washed with permeabilisation buffer. Cells were incubated with intracellular antibodies against FoxP3, Ki-67, interferon γ (IFN-γ), and granzyme B (GzmB) in permeabilisation buffer overnight at 4 °C. After washing, cells were incubated with streptavidin in permeabilisation buffer for 30 min at 4 °C. Finally, cells were washed twice with permeabilisation buffer and resuspended in FACS buffer for acquisition. For unmixing, a combination of single-stained cells and UltraComp eBeads Plus (Invitrogen) was used to generate reference controls. Sample acquisition was performed with a 3-laser Aurora spectral flow cytometer equipped with three excitation lasers (405, 488, and 640 nm) and the enhanced small particle (ES) detector module (Cytek). Samples were acquired using SpectroFlo software (v3.3.0; Cytek). Prior to acquisition, daily quality control was performed according to the manufacturer’s instructions.

**Table 1.**
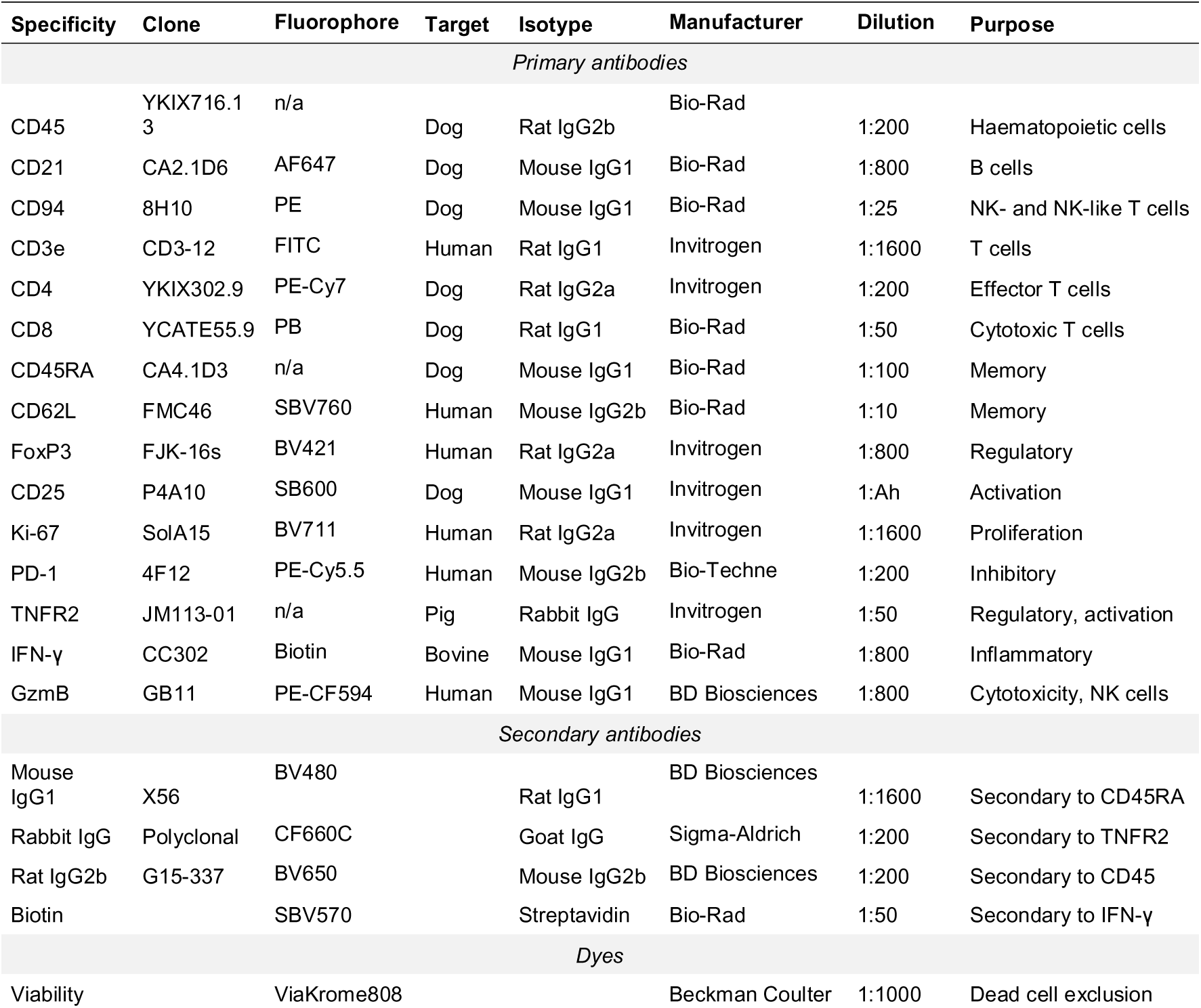
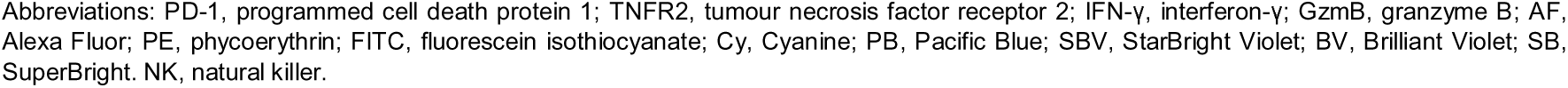
Antibodies and dyes included in the canine T cell panel.

| Specificity | Clone | Fluorophore | Target | Isotype | Manufacturer | Dilution | Purpose |
| --- | --- | --- | --- | --- | --- | --- | --- |
| *Primary antibodies* |  |  |  |  |  |  |  |
| CD45 | YKIX716.1<br>3 | n/a | Dog | Rat IgG2b | Bio-Rad | 1:200 | Haematopoietic cells |
| CD21 | CA2.1D6 | AF647 | Dog | Mouse IgG1 | Bio-Rad | 1:800 | B cells |
| CD94 | 8H10 | PE | Dog | Mouse IgG1 | Bio-Rad | 1:25 | NK- and NK-like T cells |
| CD3e | CD3-12 | FITC | Human | Rat IgG1 | Invitrogen | 1:1600 | T cells |
| CD4 | YKIX302.9 | PE-Cy7 | Dog | Rat IgG2a | Invitrogen | 1:200 | Effector T cells |
| CD8 | YCATE55.9 | PB | Dog | Rat IgG1 | Bio-Rad | 1:50 | Cytotoxic T cells |
| CD45RA | CA4.1D3 | n/a | Dog | Mouse IgG1 | Bio-Rad | 1:100 | Memory |
| CD62L | FMC46 | SBV760 | Human | Mouse IgG2b | Bio-Rad | 1:10 | Memory |
| FoxP3 | FJK-16s | BV421 | Human | Rat IgG2a | Invitrogen | 1:800 | Regulatory |
| CD25 | P4A10 | SB600 | Dog | Mouse IgG1 | Invitrogen | 1:Ah | Activation |
| Ki-67 | SoIA15 | BV711 | Human | Rat IgG2a | Invitrogen | 1:1600 | Proliferation |
| PD-1 | 4F12 | PE-Cy5.5 | Human | Mouse IgG2b | Bio-Techne | 1:200 | Inhibitory |
| TNFR2 | JM113-01 | n/a | Pig | Rabbit IgG | Invitrogen | 1:50 | Regulatory, activation |
| IFN-γ | CC302 | Biotin | Bovine | Mouse IgG1 | Bio-Rad | 1:800 | Inflammatory |
| GzmB | GB11 | PE-CF594 | Human | Mouse IgG1 | BD Biosciences | 1:800 | Cytotoxicity, NK cells |
| *Secondary antibodies* |  |  |  |  |  |  |  |
| Mouse IgG1 | X56 | BV480 |  | Rat IgG1 | BD Biosciences | 1:1600 | Secondary to CD45RA |
| Rabbit IgG | Polyclonal | CF660C |  | Goat IgG | Sigma-Aldrich | 1:200 | Secondary to TNFR2 |
| Rat IgG2b | G15-337 | BV650 |  | Mouse IgG2b | BD Biosciences | 1:200 | Secondary to CD45 |
| Biotin |  | SBV570 |  | Streptavidin | Bio-Rad | 1:50 | Secondary to IFN-γ |
| *Dyes* |  |  |  |  |  |  |  |
| Viability |  | ViaKrome808 |  |  | Beckman Coulter | 1:1000 | Dead cell exclusion |

Abbreviations: PD-1, programmed cell death protein 1; TNFR2, tumour necrosis factor receptor 2; IFN-γ , interferon- $\gamma$ ; GzmB, granzyme B; AF, Alexa Fluor; PE, phycoerythrin; FITC, fluorescein isothiocyanate; Cy, Cyanine; PB, Pacific Blue; SBV, StarBright Violet; BV, Brilliant Violet; SB, SuperBright. NK, natural killer.

### 2.5 Data analysis

Data analysis was done using SpectroFlo (v3.3.0) to generate and apply unmixing matrices, followed by FlowJo (v10.10; BD Biosciences), both via manual gating and t-distributed stochastic neighbour embedding (t-SNE) cluster analysis. For the t-SNE cluster analysis, samples with a minimum of 5,000 T cells were included to ensure sufficient resolution for the detection of rare T cell populations. Data were downsampled to include an equal number of events and then concatenated for further analysis. T cell clusters were identified and manually gated in the t-SNE map based on the signal intensity of the phenotypic markers.

Heatmaps of the Median Fluorescent Intensities (MFIs) of the identified clusters were made in R Studio (v4.4.0). Graphs and statistical analyses were performed using GraphPad Prism (v10.4.1). Statistical comparisons between media and PHA-stimulated conditions were made using a paired t-test. Statistical comparisons between tumour-infiltrating, tissue-resident, and circulating lymphocytes were made using an ordinary one-way ANOVA, while comparisons between tumour-infiltrating and circulating lymphocytes were performed using unpaired t-tests.

## 3 Results

### 3.1 Study population

A total of 11 tumour samples were collected, comprising oral melanomas (*n* = 8), an oral mast cell tumour (MCT, *n =1*), an oral peripheral nerve sheath tumour (PNST, *n =1*) and an adrenal cortical carcinoma (ACC, *n =1*). Diagnoses were obtained from the referring veterinarian and based on histopathological evaluation. Seven melanomas were recurrent, with two originating from the same patient. Patients ranged in age from 6 to 15 years old with a male-to-female ratio of 7:4 and represented different breeds, including a Great Dane, Golden Retriever, Labrador Retriever, Jack Russel Terrier, Parson Russel Terrier, Airedale Terrier, Boomer, English Cocker Spaniel, Shih Tzu, and mixed breed.

### 3.2 Mitogen stimulation induces activated and exhausted T cell phenotypes

We developed a spectral flow cytometry panel for comprehensive T cell characterisation, enabling simultaneous assessment of T cell phenotype and function. To examine panel performance and T cell mitogen-induced changes in T cell phenotype, canine PBMC from five healthy donors were cultured for 72 h in the presence or absence of PHA and then stained with the selected antibody panel (Table 1, Figures 1, S1, S2). Gating strategy is depicted in Figure S1A. First, we analysed changes in immune cell subsets. As shown in Figure S2A, upon stimulation, CD8^+^ T cell frequencies increased amongst CD3^+^ T cells, while CD4^+^ T cell frequencies decreased in all donors but one. CD4^-^CD8^-^ T cell frequencies also decreased (Figure S2A). T cell memory subsets were characterised using CD45RA and CD62L, identifying naïve T cells (T_N_; CD62L^+^CD45RA^+^), central memory T cells (T_CM_; CD62L^+^CD45RA^-^), effector memory T cells (T_EM_; CD62L^-^CD45RA^-^), and terminally differentiated memory T cells (T_EMRA_; CD62L^-^CD45RA^+^) (Bauman et al., 2022; Novak et al., 2023; Withers et al., 2018). Generally, we observed an increase in T cell differentiation (Figure 1A-C), as demonstrated by a decrease in the fraction of CD8^+^ T_N_, CD4^+^ and CD8^+^ T_CM_, and CD4^+^ T_EM_, and an increase in the fraction of CD4^+^ and CD8^+^ T_EMRA_ (Figure 1B and C). The fraction of CD4^+^ T_N_ slightly decreased in all donors except one, although this change did not reach statistical significance. Furthermore, B cell frequencies decreased upon T cell mitogen stimulation (Figure S2B). Finally, CD25^+^FoxP3^+^ T_regs_ could be detected in unstimulated PBMC; however, mitogen-induced upregulation of both markers hampered T_reg_ identification after stimulation (Figure S1B).

**Figure 1.**
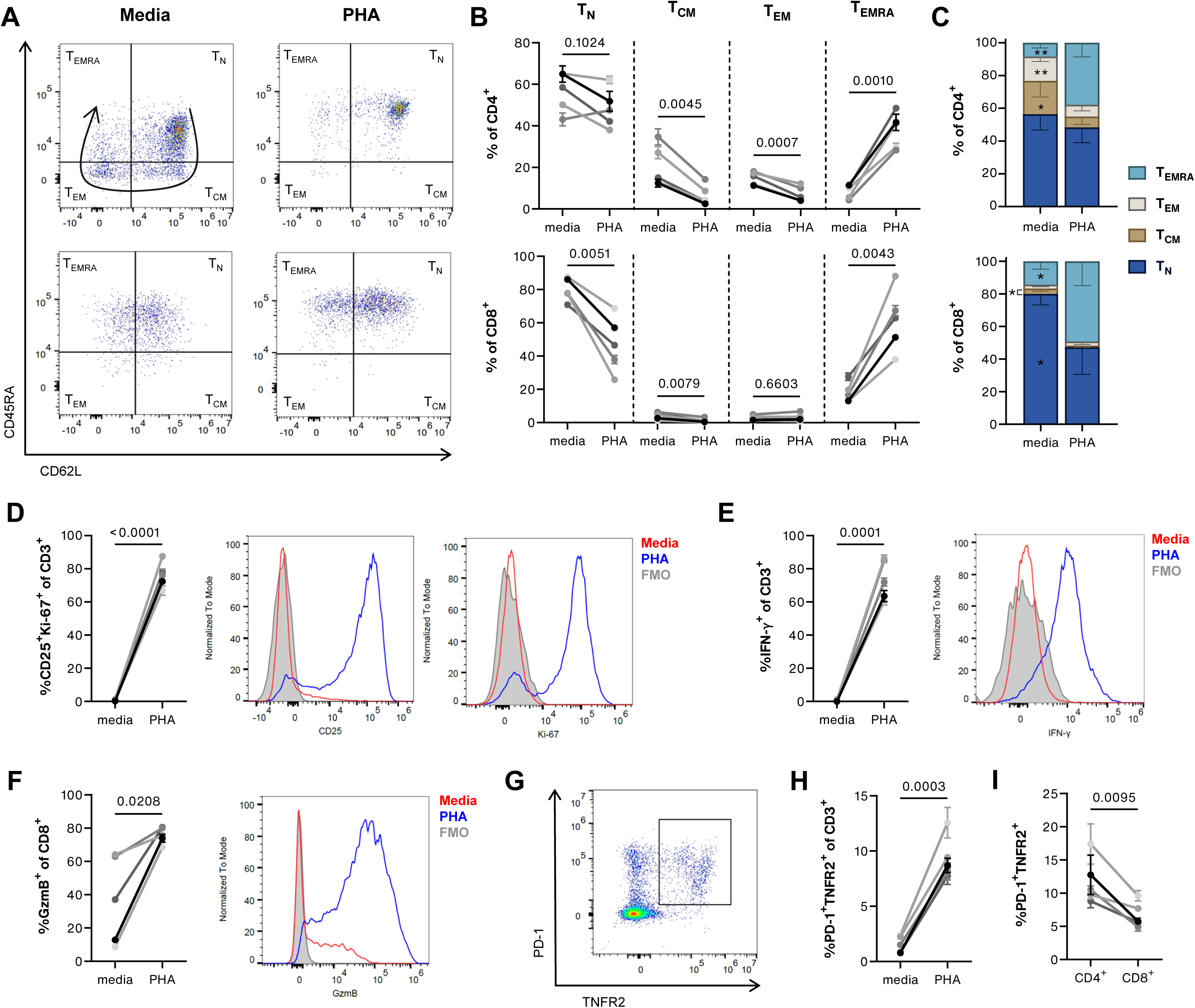
Mitogen-induced changes in T cell phenotypes. Canine PBMC (*n* = 5) were cultured in triplicate for 72 h in the presence or absence of 1 μg/ml PHA and analysed with spectral flow cytometry. (A) Flow cytometry plots of live, single, memory CD4^+^ and CD8^+^ T cell subsets, characterising naïve T cells (T_N_; CD62L^+^CD45RA^+^), central memory T cells (T_CM_; CD62L^+^CD45RA^-^), effector memory T cells (T_EM_; CD62L^-^CD45RA^-^), and terminally differentiated effector memory T cells (T_EMRA_; CD62L^-^CD45RA^+^). The arrow represents the differentiation trajectory. (B and C) Mitogen-induced changes in memory subsets as a percentage of CD4^+^ or CD8^+^ T cells, depicted in dot plots (B) or stacked bar charts (C). (D-H) Mitogen-induced proportional changes in functional markers CD25 and Ki-67 (D), IFN-γ (E), granzyme B (F), PD-1 and TNFR2 (G-I). (B-F) Histogram plots include fluorescence minus one (FMO) controls. (G) Flow cytometry plot of PHA-stimulated live, single CD3^+^ cells demonstrates PD-1^+^TNFR2^-^ and PD-1^+^TNFR2^+^ expressing populations. (H) The percentage of PD-1^+^TNFR2^+^ cells is increased in live, single CD3^+^ cells. (I) PHA-stimulated CD4^+^ T cells express more PD-1 and TNFR2 than CD8^+^ T cells. Values in dot plots represent *p*-values. Asterisks in bar charts resemble significant change between media and PHA-stimulated conditions; * ≤ 0.05, ** ≤ 0.01.

Next, we examined the effect of mitogen stimulation on functional markers within the different subsets. As expected, activation marker CD25 and proliferation marker Ki-67 increased in all T cell subsets (Figure 1D and S1C), more predominantly in CD4^+^ than in CD8^+^ T cells (Figure S2C). The number of IFN-γ^+^ cells was also increased in all T cell subsets (Figure 1E and S1D), and GzmB production was increased in CD8^+^ T cells (Figure 1F and S1E). Expression of inhibitory receptor PD-1 increased upon mitogen stimulation in CD4^+^ and CD8^+^ T cell subsets (Figure S1F and S2D). In both stimulated and unstimulated conditions, more PD-1 was observed on CD4^+^ than on CD8^+^ T cells (Figure S2E). Almost half of the PHA-stimulated and a quarter of the unstimulated PD-1^+^ cells also expressed TNFR2^+^ (Figure 1G and S2F). This PD-1^+^TNFR2^+^ subset also increased upon mitogen stimulation (Figure 1H and S1F), most predominantly in CD4^+^ than in CD8^+^ T cells (Figure 1I).

Finally, we performed a t-SNE cluster analysis including CD3^+^ T cells from both stimulated and unstimulated conditions for each donor (*n* = 5), with each condition assessed in triplicate (Figure 2). T_N_, T_CM_, T_EM_, and T_EMRA_ were present in all T cell subsets (CD4^+^, CD8^+^, and CD4^-^ CD8^-^), whereas PD1^+^TNFR2^+/-^ T cells were only observed within CD4^+^ and CD8^+^ T cells (Figure 2A and B). Comparing cluster proportions between the media and PHA-stimulated conditions, we observed an increase in T_EMRA_ and a decrease in T_N_, as well as an induction of PD1^+^TNFR2^+/-^ populations in both CD4^+^ and CD8^+^ T cells (Figure 2B). CD4^-^CD8^-^ T cells were rarely detected under stimulated conditions. T cell subsets present under both stimulated and unstimulated conditions (≥ 2% of total CD3^+^ T cells) included CD4^+^ and CD8^+^ T_N_ and T_EMRA_, as well as CD4^+^ PD-1^+^ T cells. In these subsets, mitogen stimulation upregulated FoxP3, CD25, Ki-67, IFN-γ, and GzmB (Figure 2C). Overall, these data demonstrate that stimulated and unstimulated T cell populations form distinct clusters (Figure 2A and D), confirming that our panel captures different canine T cell subsets.

**Figure 2.**
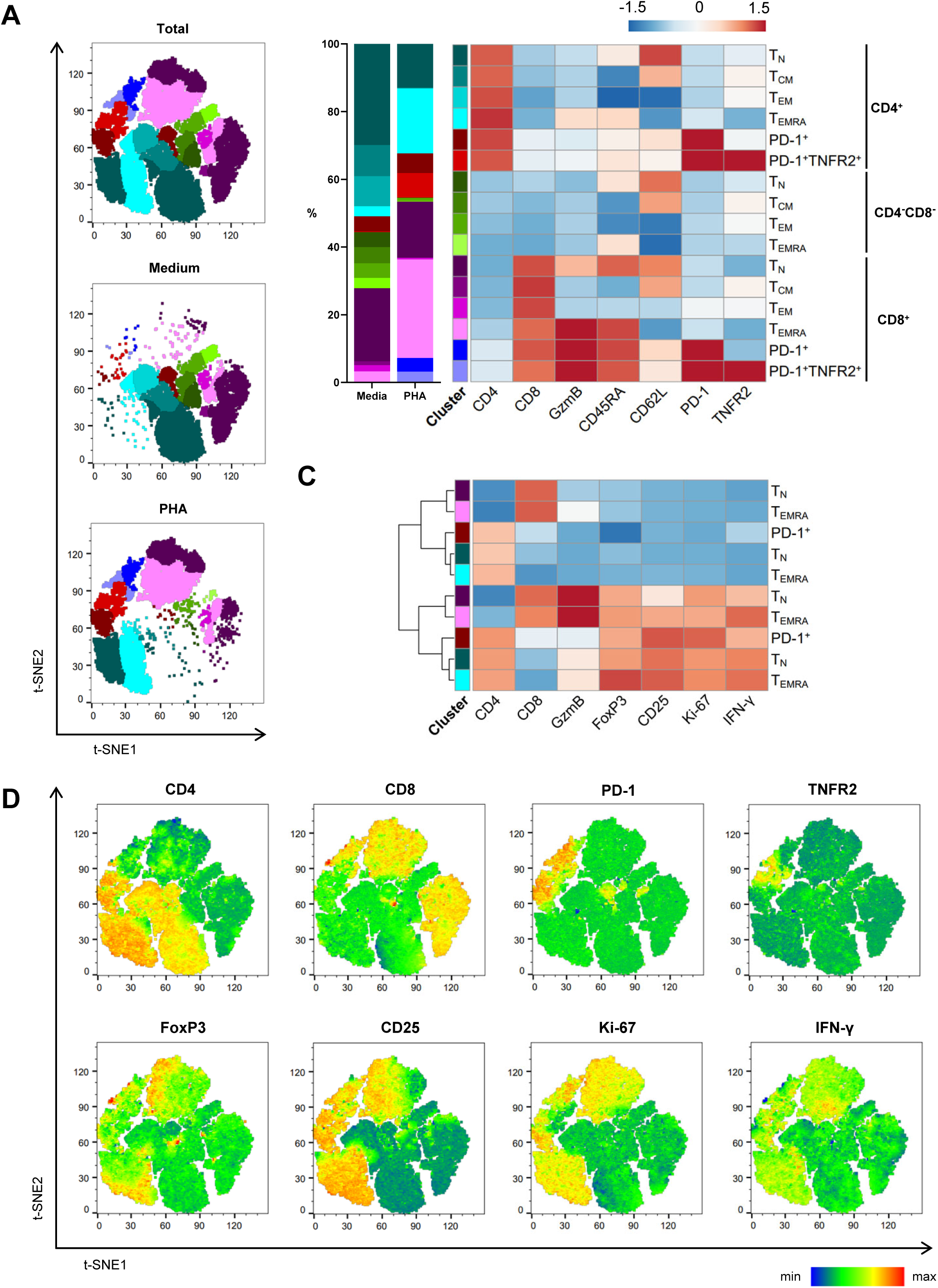
t-SNE clustering analysis of media and PHA-stimulated live, single CD3^+^ cells. Canine PBMC (*n* = 5) were cultured in triplicate for 72 h in the presence or absence of 1 μg/ml PHA and analysed with spectral flow cytometry. (A) Flowcytometry plots of T cell clusters, illustrating cluster distribution in total events, in media or in PHA-stimulated conditions. (B) Heatmap showing Median Fluorescent Intensities (MFIs) of lineage markers within each T cell subset. The proportion of each subset in media or PHA-stimulated conditions is represented in stacked bar charts. (C) Heatmap showing MFIs of functional markers in T cell subsets shared between stimulated and unstimulated conditions. (D) Flowcytometry heatmap plots showing MFI of CD4, CD8, PD-1, TNFR2, FoxP3, CD25, Ki-67 and IFN-γ. Relatively, red resembles a high MFI and blue a low MFI.

### 3.3 Higher cell yield and CD45^+^ cell frequency in solid tumours than in oral mucosa biopsies

To enable immunophenotypic analysis of tumour-infiltrating and tissue-resident immunophenotypes, biopsies were enzymatically digested to generate a single-cell suspension. The number of cells obtained from tumour biopsies (*n* = 11) was higher than that from oral mucosa biopsies (*n =* 4). As shown in Figure S3A, tumour biopsies yielded an average of 14 million cells per gram (range: 0.02–60 million), whereas oral mucosa biopsies yielded an average of 0.5 million cells per gram (range: 0.25–0.67 million). The gating strategy used to determine CD45^+^ and CD3^+^ populations in the single-cell suspensions is shown in Figure S3D. The frequency of CD45^+^ cells was higher in the tumour than in the oral mucosa samples (13% versus 5%), although the difference did not reach statistical significance (*p* = 0.1422; Figure S3B). This was due to the substantial variation in the percentage of CD45^+^ cells amongst tumour samples (range: 1.5–21%). In contrast, the frequency of CD3^+^ cells did not differ between tumour and oral mucosa samples (1.5% versus 2%; Figure S3C).

### 3.4 Canine oral malignant melanoma demonstrates a protumour immune microenvironment

To characterise the TIL phenotypes in COMM, we applied our panel to COMM single-cell suspensions (*n = 8*) and compared the T cell phenotypes with those in the oral mucosa (*n = 4*) and blood (*n = 8*) from healthy donors. Gating strategy is depicted in Figure S4. CD94 was added to the panel to serve as an NK (CD3^-^CD94^+^GzmB^+^) and NKT (CD3^+^CD94^+^) cell marker(Graves et al., 2019; Scorza et al., 2022). NKT and NK cell frequencies were decreased, although the latter did not reach significance (*p* = 0.779) (Figure 3A and S5A). We observed that TILs had higher CD4^+^ and lower CD8^+^ T cell frequencies than tissue-resident and circulating lymphocytes, resulting in an elevated CD4:CD8 ratio (Figure 3B and S5B-D). Regarding memory subsets, COMM contained a higher proportion of T_EM_, and lower proportions of T_N_, CD4^+^ and CD4^-^CD8^-^ T_CM_, and CD4^+^ and CD8^+^ T_EMRA_ relative to the blood (Figure 3C). In comparison with tissue-resident lymphocytes, TILs displayed relatively fewer CD4^-^CD8^-^ T_EMRA_. Furthermore, a larger proportion of CD4^+^ TILs were CD25^+^FoxP3^+^ compared to circulating CD4^+^ T cells (Figure 3D). A similar trend was observed between CD25^+^FoxP3^+^ CD4^+^ TILs and tissue-resident CD4^+^ T cells. CD25^+^FoxP3^+^ cells were also detected within the CD4^-^CD8^-^ T cell subset, but their frequency did not differ significantly across tissues (Figure S5E). Most CD25^+^FoxP3^+^ cells displayed a T_EM_ phenotype (Figure S4D and S5F). Finally, the B cell proportion in COMM and the oral mucosa was less than in the blood (Figure S5G).

**Figure 3.**
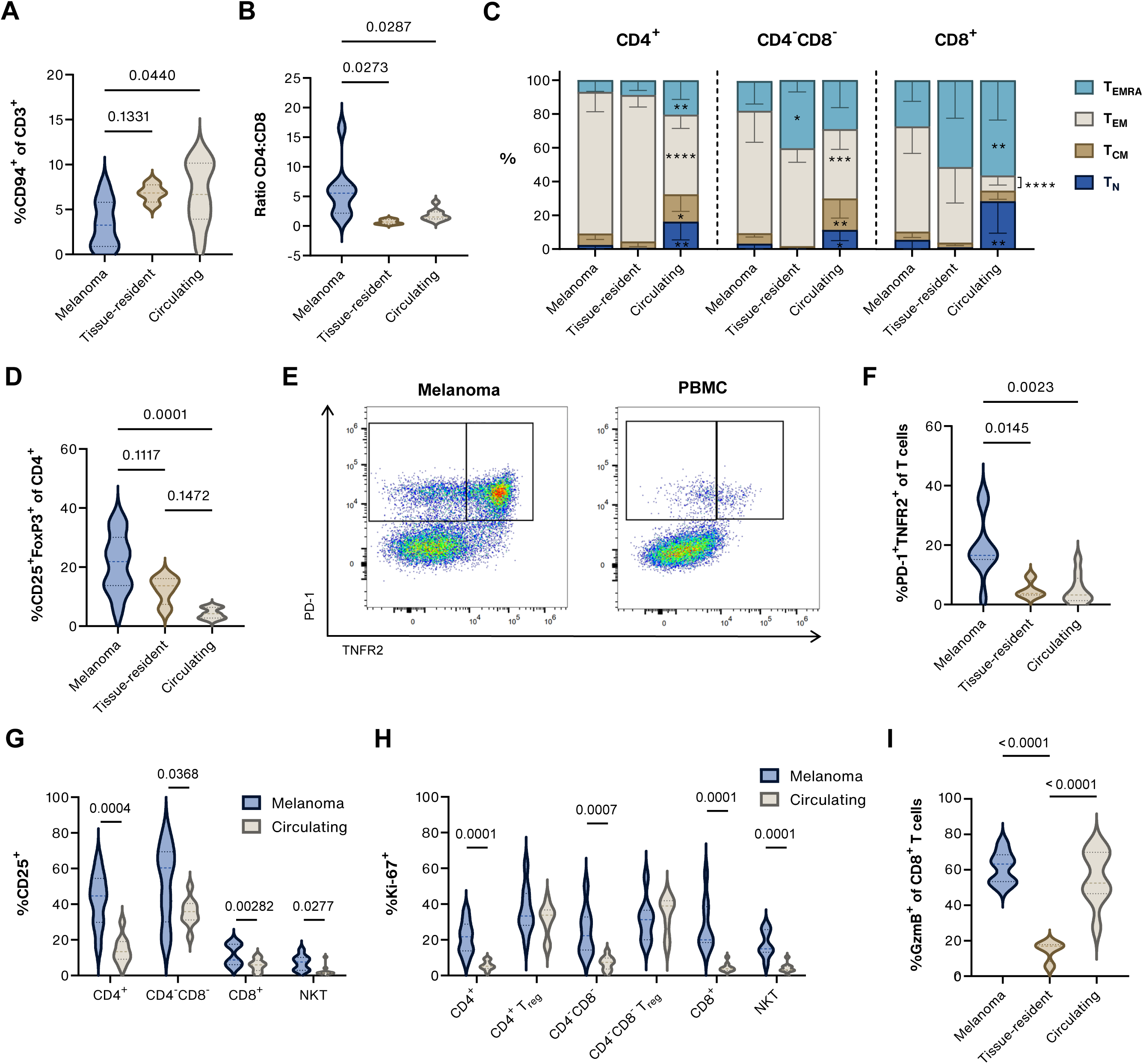
Characterisation of different tumour-infiltrating, tissue-resident and circulating immunophenotypes. Single-cell suspension of melanoma samples (*n* = 5), healthy oral mucosa (*n* = 4), and healthy PBMC (*n* = 5) were analysed with spectral flow cytometry. (A-D) Proportion of CD3^+^CD94^+^ NKT cells (A), CD4:CD8 ratio (B), CD4^+^ and CD8^+^ memory subsets (C), and CD25^+^FoxP3^+^ cells (D) in the different tissues. Memory subsets were characterised as naïve T cells (T_N_; CD62L^+^CD45RA^+^), central memory T cells (T_CM_; CD62L^+^CD45RA^-^), effector memory T cells (T_EM_; CD62L^-^CD45RA^-^), and terminally differentiated effector memory T cells (T_EMRA_; CD62L^-^CD45RA^+^). (E-H) Expression of PD-1 and TNFR2 (E, F), CD25 (G), Ki-67 (H), and granzyme B (I) in tumour-infiltrating, tissue-resident, and circulating lymphocytes. CD4^+^ and CD4^-^CD8^-^ T cells were defined as non-Tregs. Values in bar charts represent p-values. Asterisks in bar charts resemble significant change with melanoma TILs; * ≤ 0.05, ** ≤ 0.01, *** ≤ 0.001, **** ≤ 0.0001.

Next, we assessed functional markers within the different T cell subsets. Importantly, we detected significantly higher proportions of PD1^+^TNFR2^+/-^ cells in all TIL subsets than in the tissue-resident or circulating T cell subsets (Figure 3E, F and S5H). Relative to the blood, COMM samples contained higher proportions of CD4^+^, CD4^-^CD8^-^, CD8^+^, and NKT cells expressing either activation marker CD25 or proliferation marker Ki-67 (Figure 3G and H). Similarly, CD25^+^Ki-67^+^ cell frequencies were significantly increased in these subsets (Figure S5I). Finally, a similar percentage of tumour-infiltrating and circulating NKT and CD8^+^ T cells was GzmB^+^, whereas tissue-resident CD8^+^ T cells had lower levels of GzmB (Figure 3I). This comparison could not be made for tissue-resident NKT cells due to the low number of detected events, although pooling the oral mucosa samples revealed a much lower percentage of tissue-resident GzmB^+^ NKT cells than NKT TILs (29% versus 86%; Figure S5J).

To validate the protumour TME in canine melanoma identified through manual gating, we performed a t-SNE cluster analysis. Three melanoma samples and the healthy oral mucosa samples were excluded from the analysis because they contained fewer than 5,000 T cells. Five PBMC samples acquired on the same day as the remaining melanoma samples were included as controls. A total of 7,000 live, single T cells per sample could be used for downstream analysis. The t-SNE cluster analysis of melanoma and blood samples confirmed the enrichment of CD25^+^FoxP3^+^ CD4^+^ T_regs_, with or without expression of PD-1 and TNFR2 (Figure 4). These T_regs_ also expressed Ki-67 (Figure 4B). Despite a smaller proportion of CD8^+^ TILs, the T_reg_ frequency appears to be the main driver of the aforementioned increased CD4:CD8 TIL ratio (Figure 3B). Furthermore, we observed a smaller proportion of NKT TILs compared to circulating lymphocytes, and all expressed PD-1 (Figure 4B). Regarding memory status, T_EM_ were enriched in COMM, and a substantial proportion of tumour-infiltrating T_EM_ co-expressed PD-1 and TNFR2 (Figure 4A-B). This was most predominant in CD4^+^ TILs, but could also be observed in CD4^-^CD8^-^ and CD8^+^ TILs (Figure 4B). Compared to normal PD-1^-^TNFR2^-^ T_EM_, PD-1^+^TNFR2^+^ T_EM_ upregulated Ki-67 in all TIL subsets (Figure 4B). Additionally, CD4^-^CD8^-^ T_EM_ upregulated CD25, whereas CD8^+^ T_EM_ upregulated GzmB (Figure 4B). Generally, we observed in all T cell subsets relatively fewer T_EMRA_ in COMM than in blood (Figure 4A-B). In contrast to T_EM_, a small proportion of T_EMRA_ expressed PD-1 but not TNFR2. These PD-1^+^ T_EMRA_ lacked Ki-67 expression, and PD-1^+^ CD8^+^ T_EMRA_ lacked GzmB (Figure 4B). TILs hardly expressed the lymph node homing marker CD62L, and these T_N_ and T_CM_ did not express PD-1 or TNFR2 (Figure 4B). Together, these findings indicate a tumour-promoting TME in COMM, characterised by diminished cytotoxic capacity and enrichment of regulatory as well as activated and exhausted phenotypes.

**Figure 4.**
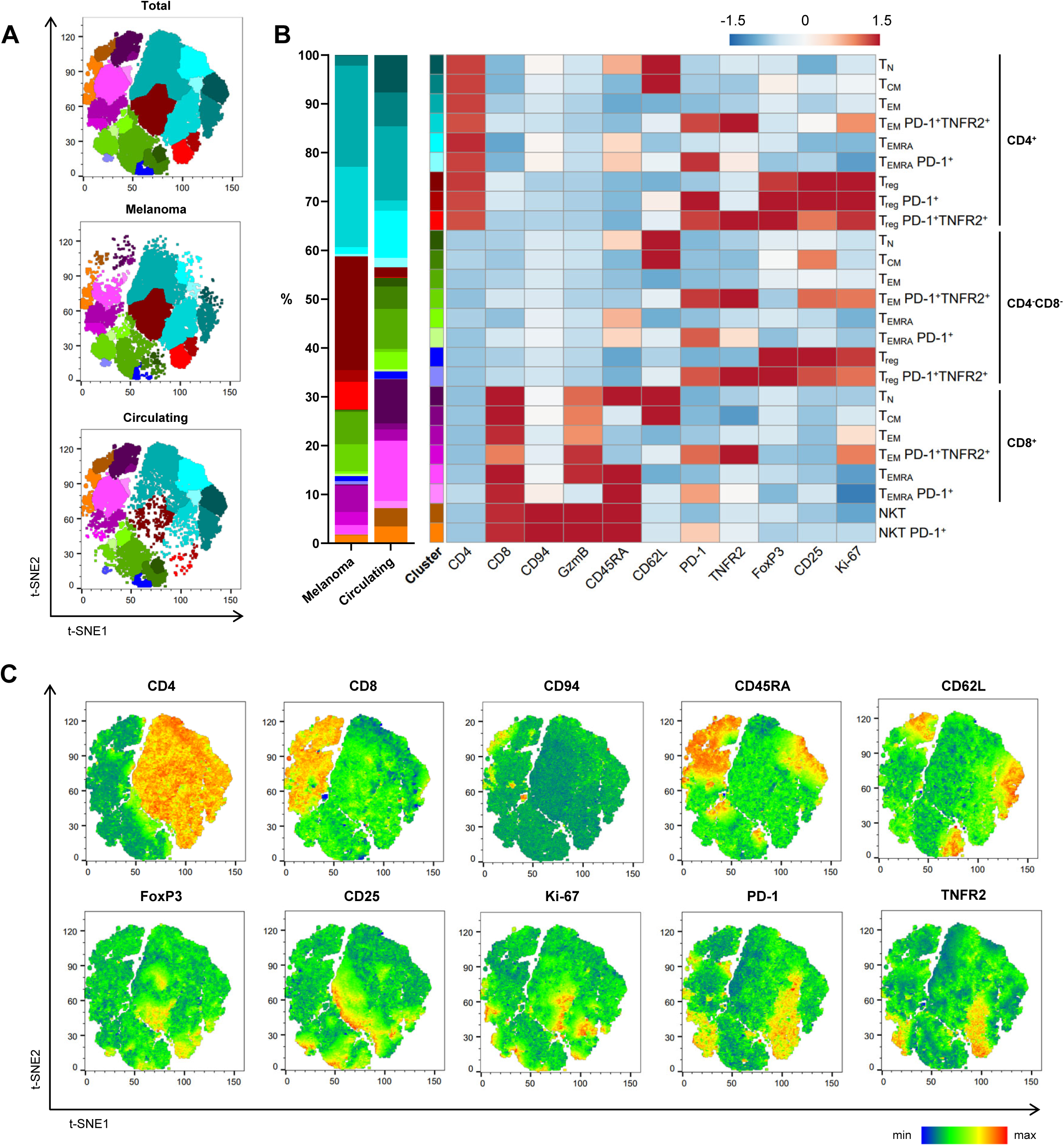
t-SNE cluster analysis of tumour-infiltrating and circulating live, single CD3^+^ cells. Single-cell suspension of melanoma samples (*n* = 5) and healthy PBMC (*n* = 5) were analysed with spectral flow cytometry. (A) Flow cytometry plots of T cell clusters, illustrating cluster distribution in total events, in melanoma or in blood. (B) Heatmap showing Median Fluorescent Intensity (MFIs) of the markers within each cluster. The frequency of each cluster is represented in stacked bar charts. (C) Flow cytometry heatmaps showing MFI of CD4, CD8, CD94, CD45RA, CD62L, FoxP3, CD25, Ki-67, PD-1 and TNFR2. Relatively, red resembles a high MFI and blue a low MFI.

### 3.5 TIL phenotype in other solid tumours

To evaluate whether our panel is applicable to other solid tumours and to explore potential shared characteristics of TILs, we additionally analysed single-cell suspensions from an oral MCT, an oral PNST and an ACC. Most importantly, a substantial proportion of TILs co-expressed PD-1 and TNFR2, in contrast to tissue-resident and circulating lymphocytes (Figure 5A). We also identified CD25^+^FoxP3^+^ CD4^+^ T_regs_ (Figure 5B). Overall, T_regs_ were more abundant in tumour tissue than in healthy oral mucosa or blood, except for the PNST. Regarding memory subsets, we confirm that T_EM_ were enriched in tumours relative to blood (Figure 5C). In general, activation and proliferation markers CD25 and Ki-67 were upregulated on TILs compared to tissue-resident and circulating lymphocytes, except on CD4^+^ ACC TILs (Figure 5D). Together, this demonstrates our panel reliably captures phenotypic and functional features of TILs across different solid tumours.

**Figure 5.**
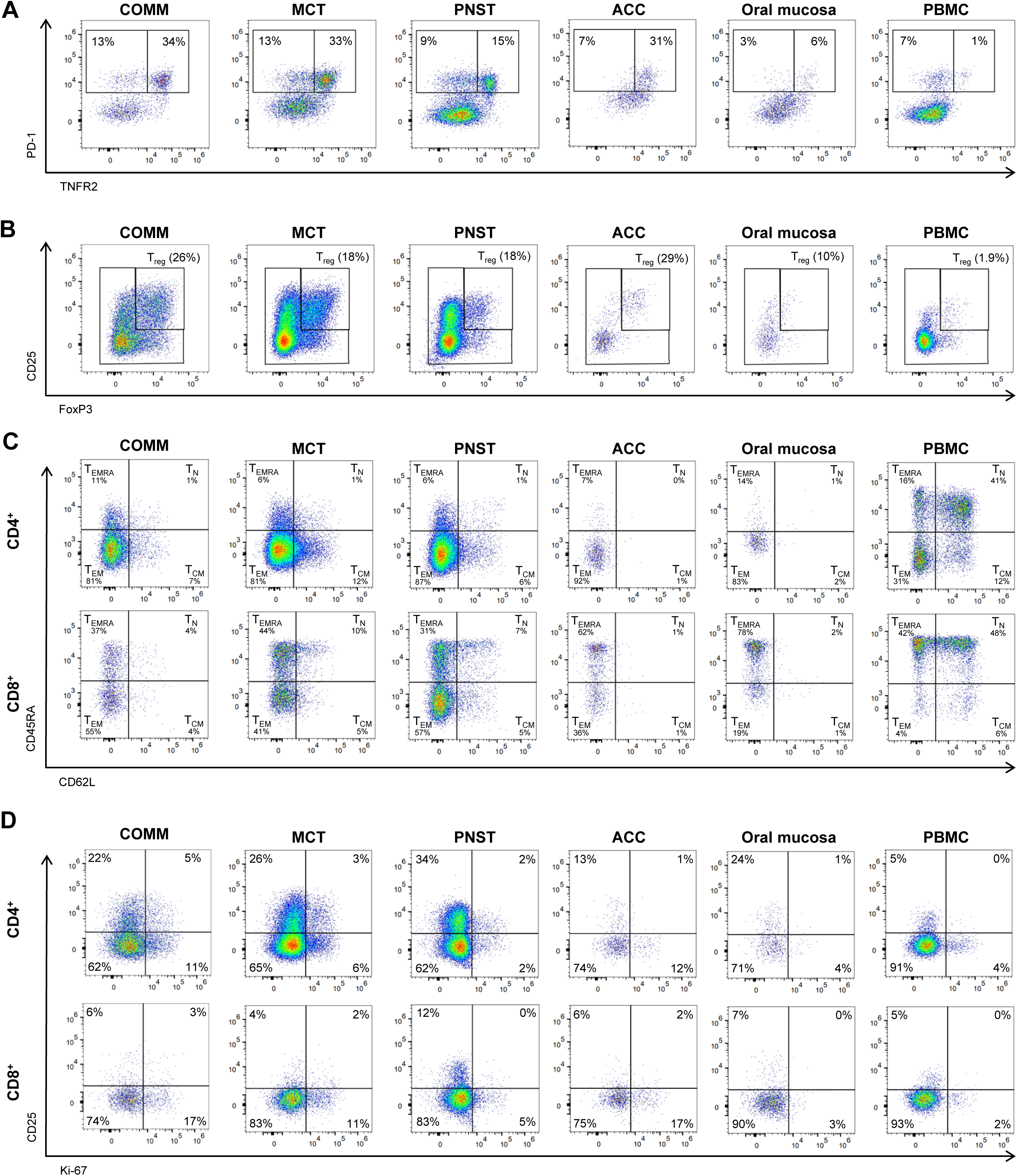
Tumour-infiltrating phenotypes in canine solid tumours. (A) PD-1 and TNFR2 in CD8^+^ T cells. (B) CD25^+^FoxP3^+^ CD4^+^ T_regs_. (C) Memory markers CD45RA and CD62L in CD4^+^ and CD8^+^ T cells. (D) Activation and proliferation markers CD25 and Ki-67 in CD4^+^ and CD8^+^ T cells. CD4^+^ represent non-T_regs_. Percentages represent frequent of parent. COMM, canine oral malignant melanoma; MCT, mast cell tumour; PNST, peripheral nerve sheath tumour; ACC, adrenal cortical carcinoma; PBMC, peripheral blood mononuclear cells.

## 4 Discussion

Extensive immunophenotyping of TILs has enhanced our understanding of tumour-immune interactions, enabled comprehensive profiling of biomarkers and immune targets, and improved the prediction of treatment outcomes. These insights are largely derived from studies in humans rather than dogs. To date, a 9-colour flow cytometry panel represents the most extensive panel used to distinguish T cells, B cells, neutrophils, and monocytes in canine melanoma (Parys et al., 2023). Here, we introduce a 16-colour spectral flow cytometry panel to analyse TIL phenotypic and functional profiles. Using this panel, we demonstrated an immunosuppressive, tumour-promoting TME in COMM, marked by fewer cytotoxic cells, a larger proportion of T_regs_, and tumour-specific activated and exhausted T cells. We further applied this panel to determine marker dynamics in a 72 h PBMC culture with or without mitogen stimulation. Overall, this study expands our understanding of canine TIL phenotype and function, helping to bridge the knowledge gaps that currently limit comparative oncology approaches.

CD8^+^ T cells, NKT and NK cells can directly kill tumour cells, and their abundance is essential in antitumour immunity (Iyoda et al., 2023; Kraja et al., 2025). In dogs, a true NK cell marker is lacking, but several studies indicate that CD94 is exclusively expressed on canine NK and NKT cells (Graves et al., 2019; Scorza et al., 2022). We observed relatively few CD8^+^ T cells and NKT cells (CD3^+^CD94^+^CD8^+^) in COMM compared to the oral mucosa or blood. However, based on GzmB expression, CD8^+^ TILs were significantly more cytotoxic than tissue-resident, but not circulating CD8^+^ T cells, demonstrating compartmental differences in immune cell phenotypical and functional profiles. Consistent with this, others report similar cytotoxicity in tumour-infiltrating and circulating CD8^+^ T cells from both healthy and melanoma-bearing dogs, although in the absence of significant differences in CD8^+^ T cell frequencies (Sparger et al., 2021).

T_regs_ actively suppress the immune response and have been associated with an impaired antitumour immunity (Kraja et al., 2025). Others previously observed higher frequencies of FoxP3^+^ T_regs_ in COMM relative to patient or healthy blood (Sparger et al., 2021; Tominaga et al., 2010). Consistent with this, we observed relatively more CD25^+^FoxP3^+^ T_regs_ in COMM than in healthy blood. The accumulation of T_regs_ in the oral mucosa and other barrier sites is expected, as they are essential for immunotolerance to constant antigenic stimulation (Traxinger et al., 2022). However, we observed a trend towards increased proportions of CD25^+^FoxP3^+^ T_regs_ in COMM compared to the oral mucosa (*p* = 0.1117), possibly indicative of tumour-associated accumulation.

Upon mitogen stimulation, we observed an upregulation of CD25 and FoxP3 in all T cell subsets, consistent with findings in both humans (Kmieciak et al., 2009) and dogs (Pinheiro et al., 2011). As canine-specific T_reg_ antibodies are lacking, such as CD127 and CD39, it remains uncertain if the CD25^+^FoxP3^+^ T cells in COMM represent true T_regs_ or activated conventional T cells. Similar to prior reports in humans (Vukmanovic-Stejic et al., 2008) and dogs (Sparger et al., 2021), CD25^+^FoxP3^+^ TILs indeed co-expressed Ki-67. While CD25^+^FoxP3^+^ CD8^+^ T cells were present in stimulated PBMC, they were absent in COMM, suggesting that the CD25^+^FoxP3^+^ CD4^+^ T cells detected in COMM are T_regs_, displaying regulatory activity. The lack of canine-specific T_reg_ antibodies also hampers the identification of compartment-specific differences in T_reg_ phenotype, as tissue-resident T_regs_ are found to be phenotypically distinct from circulating T_regs_ in humans (Traxinger et al., 2022).

Tissue-resident memory cells drive antitumour immunity and are correlated with improved survival (Gavil et al., 2024). In dogs, these memory subsets have been identified by CD45 splice variant CD45RA and lymph node homing marker CD62L (Bauman et al., 2022; Novak et al., 2023; Withers et al., 2018). Upon antigen encounter, T_N_ can differentiate into T_EM_, which migrate to the tissues to exert immediate effector functions (Bauman et al., 2022). Therefore, it is unsurprising that we observed an enrichment of T_EM_ and a reduction of T_N_ in the TME relative to circulation, consistent with reports in human cancers (Garman et al., 2023; Poschke et al., 2012). As antigen-specific memory responses in dogs are similarly marked by an increase in T_EM_ and a decrease in T_N_ (Novak et al., 2023), the observed T_EM_ in the TME may represent tumour-reactive cells. If antigenic stimulation persists, such as within the TME, T_EM_ can differentiate towards a terminally differentiated T_EMRA_, which may express senescence and exhaustion markers (Bauman et al., 2022). Therefore, we expected to observe higher frequencies of dysfunctional T_EMRA_ within the TME. Instead, we found relatively fewer T_EMRA_ and higher proportions of PD1^+^TNFR2^+^ T_EM_. This may reflect a preferential differentiation of T_EM_ towards an exhausted phenotype within the TME rather than a terminally differentiated phenotype (Giles et al., 2023; Jin et al., 2023).

CD4^-^CD8^-^ T cells constitute a distinct T cell population that can exert either tumour-promoting or tumour-inhibiting effects, depending on the tumour type and composition of the TME (Wu et al., 2022). In the present study, we demonstrate that CD4^-^CD8^-^ T cells comprise about 10–20% of tumour-infiltrating, tissue-resident and circulating T cells, consistent with other reports in dogs (Rabiger et al., 2019; Sparger et al., 2021). In contrast, CD4^-^CD8^-^ T cells constituted only 1–4% of T cells in human tumours and blood (Di Blasi et al., 2020; Fang et al., 2019; Fischer et al., 2005; Stankovic et al., 2019). Furthermore, we and others have shown that the immunophenotypes of CD4^-^CD8^-^ T cells in canine melanoma and blood largely overlap with those of CD4^+^ T cells, except for elevated CD25 expression (Rabiger et al., 2019; Sparger et al., 2021). Although immunoregulatory functions have been described for this subset (Protschka et al., 2024; Rabiger et al., 2019), we observed no differences in their frequency between tumour-infiltrating, tissue-resident, and circulating T cells. However, potential subset-specific differences may have been obscured since we were unable to distinguish between TCRαβ_⁺_ and TCRγδ_⁺_ CD4^-^CD8^-^ T cells, which are phenotypically distinct populations (Rabiger et al., 2019). Given their abundance and immunoregulatory role, further investigation is warranted to determine their function within the TME.

T cell exhaustion is linked to impaired antitumour immunity and characterised by overexpression of inhibitory checkpoint receptors, such as PD-1, which contributes to effector dysfunction under conditions of chronic antigenic stimulation (Zebley et al., 2024). In dogs, the PD-1/PD-L1 axis functions similarly, as its blockade improved IFN-γ production in circulating and tumour-infiltrating T cells (Choi et al., 2020; Coy et al., 2017; Nemoto et al., 2018). In addition, the costimulatory receptor TNFR2 is expressed on CD8^+^ T_ex_, as well as on highly immunosuppressive T_regs_ in humans, and its expression is correlated with expression of other immune checkpoints, including PD-1 (Gao et al., 2023; Liao et al., 2023). In the current study, we show that PD-1 expression is higher on all canine TIL subsets compared to tissue-resident or circulating T cells. Notably, the majority of these PD-1^+^ TILs co-expressed TNFR2, defining a PD-1^+^TNFR2^+^ T cell subset that is enriched in solid tumours. Although we observed an upregulation of both PD-1 and TNFR2 upon T cell stimulation, we hypothesise that the PD-1^+^TNFR2^+^ population is tumour-specific, as this subset was more abundant in the TME than among activated T cells (22% [range: 2–37%] versus 9% [range: 7–14%]). Stimulation-induced PD-1 upregulation on conventional T cells in humans and dogs has been described to mediate immune regulation (Choi et al., 2020; Nemoto et al., 2018). By comparison, stimulation-induced TNFR2 upregulation on conventional T cells in humans and mice has been associated with enhanced proliferation, effector functions, and resistance to T_regs_, whereas chronic TNFR2 stimulation on tumoral T_regs_ enhances their suppressive activity (Chen et al., 2010; Govindaraj et al., 2013). These distinct roles of PD-1 and TNFR2 suggest that dual targeting may enhance antitumour responses, as demonstrated in mouse cancer models where PD-1/PD-L1 blockade combined with TNFR2 agonism or antagonism outperformed single-agent therapy (Case et al., 2020; Chen et al., 2022; Tam et al., 2019; Zhang et al., 2022). Together, this data supports the presence of an exhausted-like PD-1^+^TNFR2^+^ T cell subset in canine tumours, which can potentially be targeted for reinvigoration by immunotherapies.

Collectively, these results highlight the complex interplay between immune activation, suppression, and dysfunction within the COMM microenvironment, emphasising the importance of considering both TIL phenotype and function. By delineating compartment-specific differences between tumour-infiltrating, tissue-resident, and circulating lymphocytes, we highlight the role of the immune environment in shaping T cell responses. However, most studies compare tumour-infiltrating T cells with circulating T cells due to the limited availability of healthy tissues, and we also faced challenges in obtaining sufficient cells from these tissues. Nevertheless, this work expands our understanding of canine tumour immunology and identifies potential targets for interventions aimed at restoring antitumour immune responses. Importantly, the 16-colour spectral flow cytometry panel here presented provides a robust tool for detailed immunophenotyping of canine TILs, that can be further expanded and tailored to bridge the knowledge gaps that currently limit comparative oncology approaches.

## Data availability statement

The data that support the findings of this study are available from the corresponding author upon reasonable request.

## Acknowledgements

We thank the Flow Cytometry and Cell Sorting Facility of The Faculty of Veterinary Medicine at Utrecht University for support. E.L.A. is ISAC SRL Emerging Leader (2025-2028). Furthermore, we thank the physicians, pathologists, and other staff of the participating clinics for their assistance in collecting, processing, and providing the tumour specimens used in this study.

