## Supplementary figures for "A 16-colour spectral flow cytometry panel to characterise T cell immunophenotypes in canine oral melanoma"

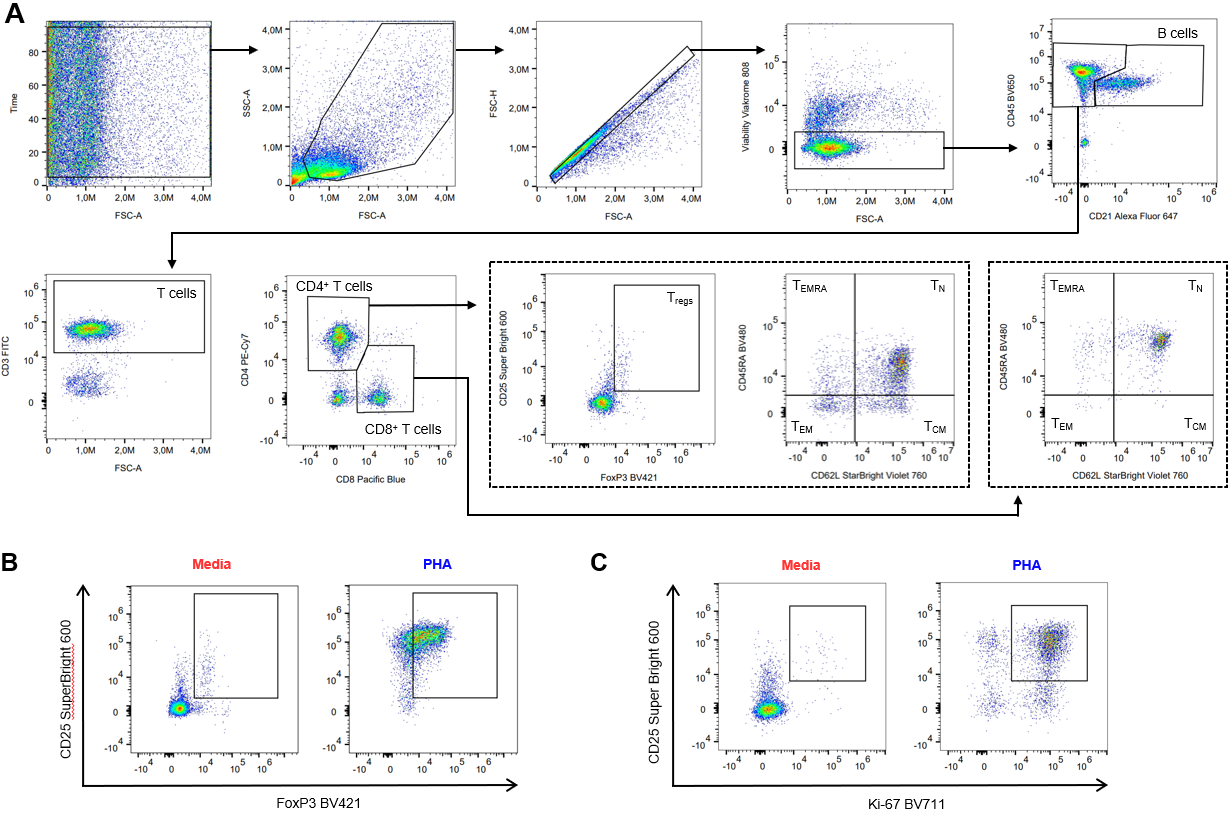

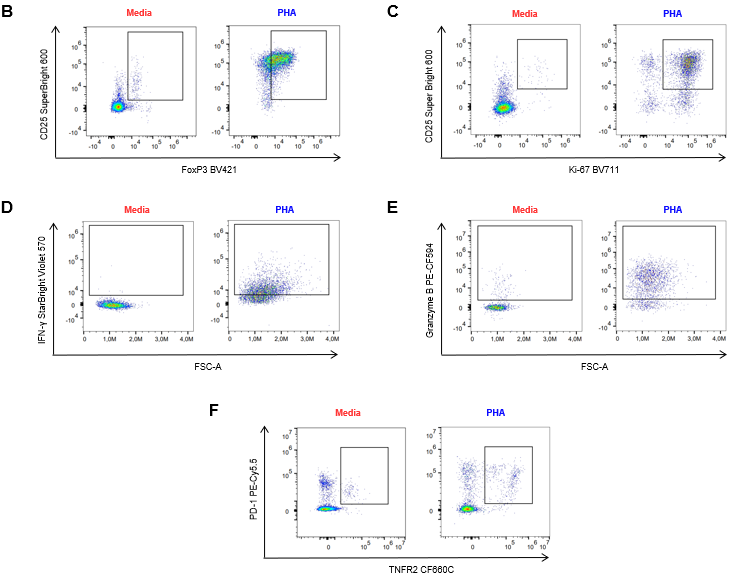

**Figure S1. Representative flow cytometry plots of canine PBMC stimulation assay.** Canine PBMC (*n* = 5) were cultured in triplicate for 72 h in the presence or absence of 1 μg/ml PHA and analysed with spectral flow cytometry. (A) Gating strategy to define immune cell subsets, including memory T cell subsets (naïve T cells, T_N_; central memory T cells, T_CM_; effector memory T cells, T_EM_; terminally differentiated memory T cells, T_EMRA_). (B) Mitogen-stimulated T cells upregulate CD25 and FoxP3, hindering regulatory T cell (T_reg_) identification. (C–E) Gating of markers within T cell subsets and their expression after mitogen stimulation: activation and proliferation marker CD25 and Ki-67 (C), IFN-γ (D), granzyme B (E), and PD-1 and TNFR2 (F).

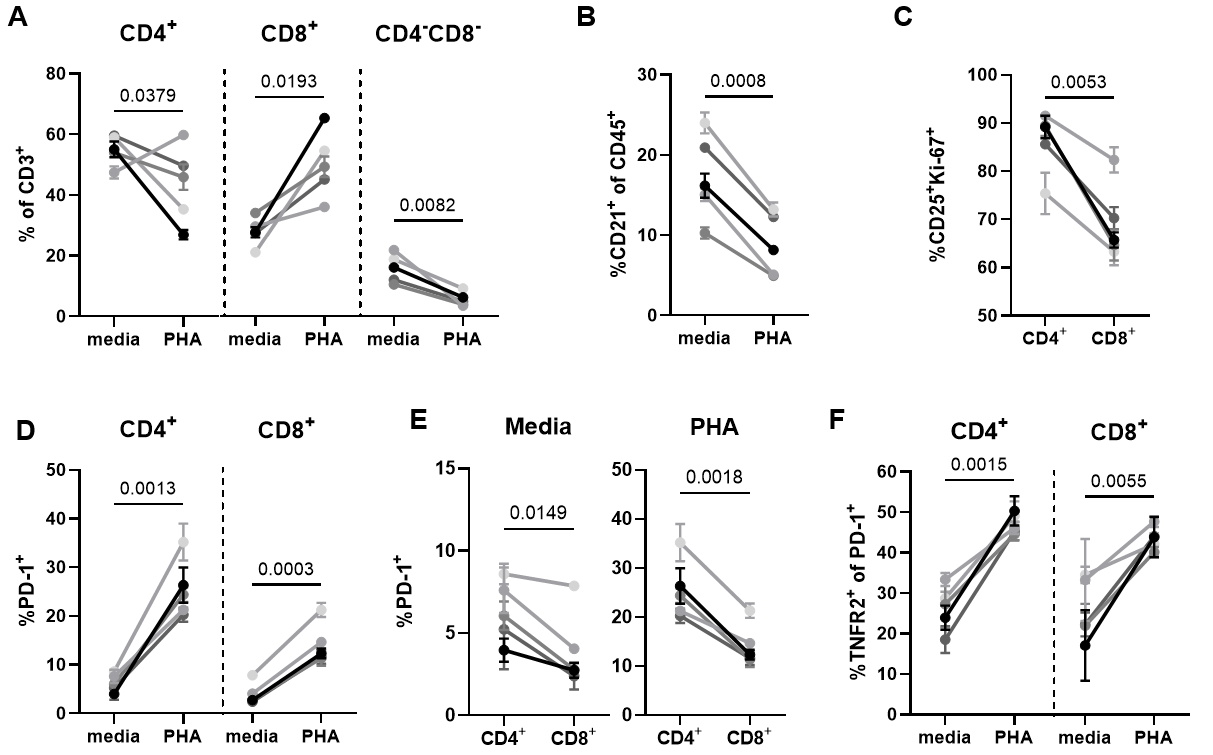

**Figure S2. Graphs of canine PBMC stimulation assay.** Canine PBMC (*n* = 5) were cultured in triplicate for 72 h in the presence or absence of 1 μg/ml PHA and analysed with spectral flow cytometry. (A) Mitogen-induced changes in CD4^+^, CD8^+^ and CD4^-^CD8^-^ subset frequencies of live, single CD3^+^ T cells. (B) Mitogen-induced changes in B cell frequencies, defined as CD21^+^ of all live, single CD45^+^ cells. (C) CD25^+^Ki-67^+^ cell frequencies in live, single PHA-stimulated CD4^+^ and CD8^+^ T cell subsets. (D) Mitogen-induced changes in PD-1^+^ frequencies in live, single CD4^+^ and CD8^+^ T cells. (E) PD-1^+^ cell frequencies in CD4^+^+ and CD8^+^ T cell subsets, either media or PHA-stimulated. (F) Mitogen-induced changes in PD-1^+^TNFR2^+^ cell frequencies in live, single CD4^+^ and CD8^+^ T cells. Lines represent individual dogs in unstimulated and stimulated conditions (A, B, D, and F) or individual dogs in either condition (B and E). Values in dot plots represent *p*-values.

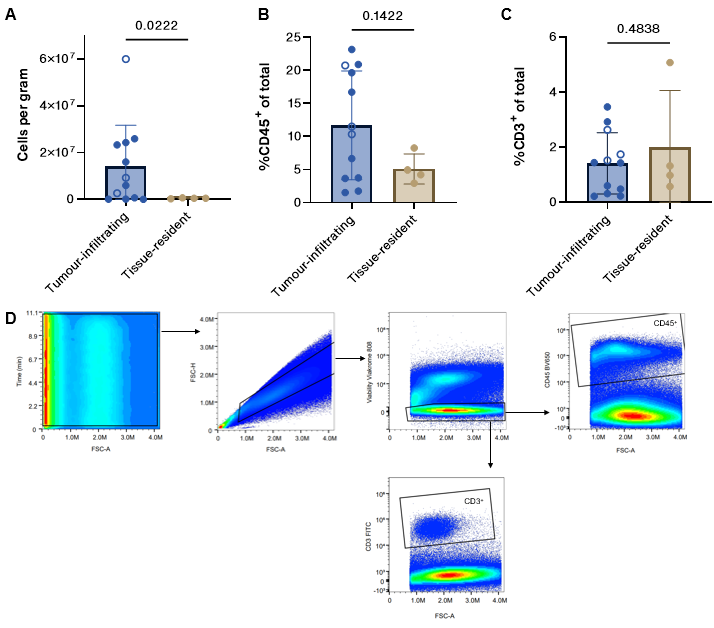

**Figure S3. Yield following tissue digestion.** (A) Number of cells per gram of tissue. (B and C) Graphs of the percentage of CD45^+^ (B) and CD3^+^ (C) cells amongst live, single cells. Open dots represent non-melanoma solid tumours: adrenal cortical carcinoma, oral mast cell tumour, and oral peripheral nerve sheath tumour. (D) Gating strategy to obtain immune cell percentages. Values in bar charts represent *p*-values.

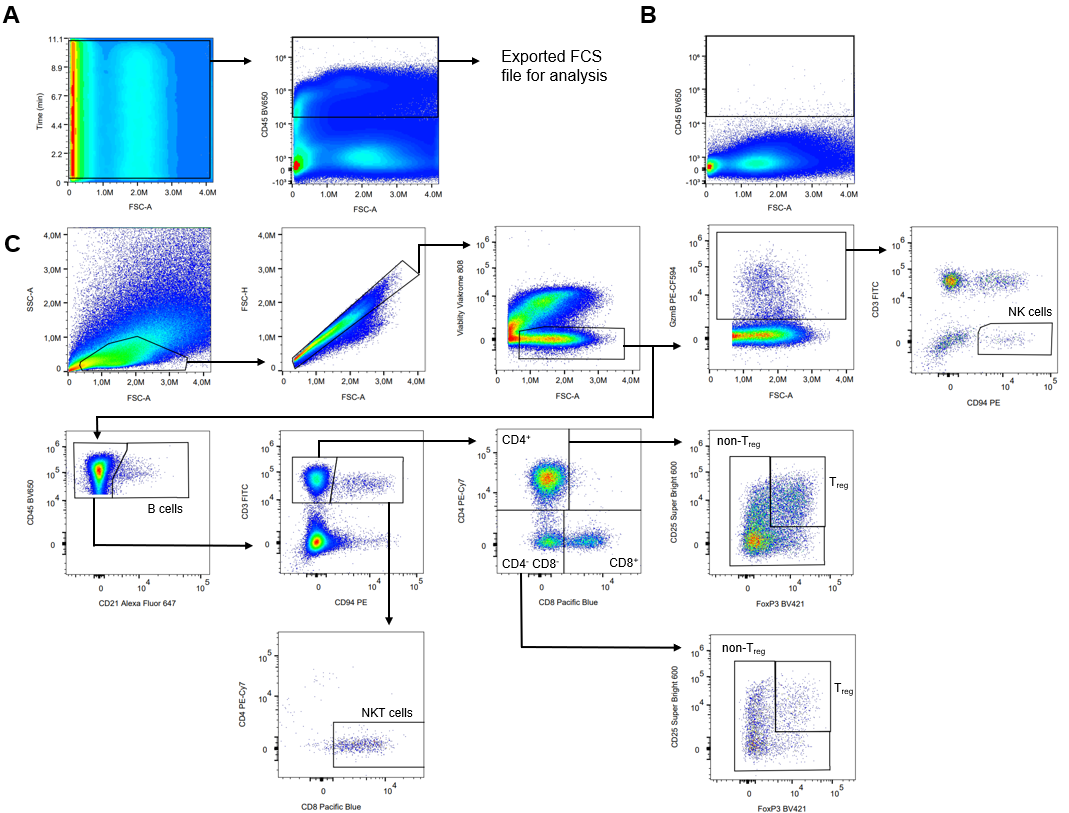

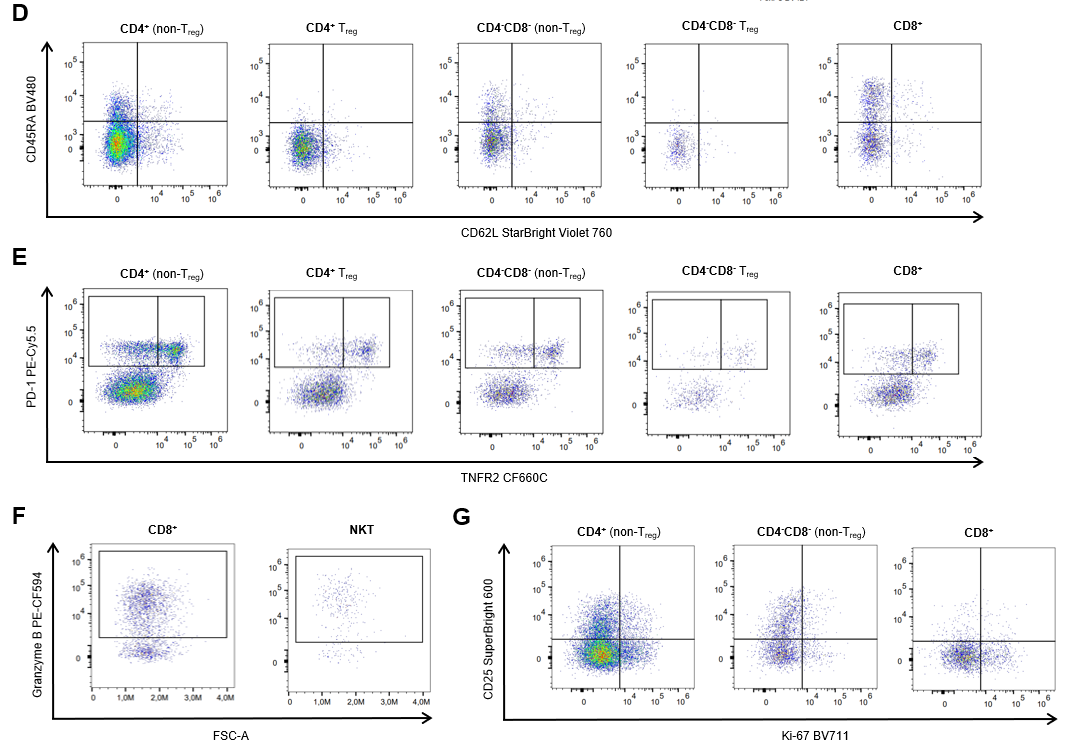

**Figure S4. Representative flow cytometry plots of melanoma single-cell suspensions.** (A and B) Gating strategy to export CD45^+^ events for data analysis (A) compared to unstained control (B). (C and D) Gating strategy to define immune cell subsets: B cells, natural killer (NK) cells, NKT cells, CD4^+^ T cells, CD8^+^ T cells, CD4^-^CD8^-^ T cells, regulatory T cells (T_regs_), and non-T_regs_ (C), as well as memory subsets, defined as CD62L^+^CD45RA^+^ naïve T cells, CD62L^+^CD45RA^-^central memory T cells, CD62L^-^CD45RA^-^ effector memory T cells, and CD62L^-^CD45RA^+^ terminally differentiated memory T cells (D). (E–G) Gating of markers within T cell subsets: PD-1 and TNFR2 (E), granzyme B (F), CD25 and Ki-67 (G).

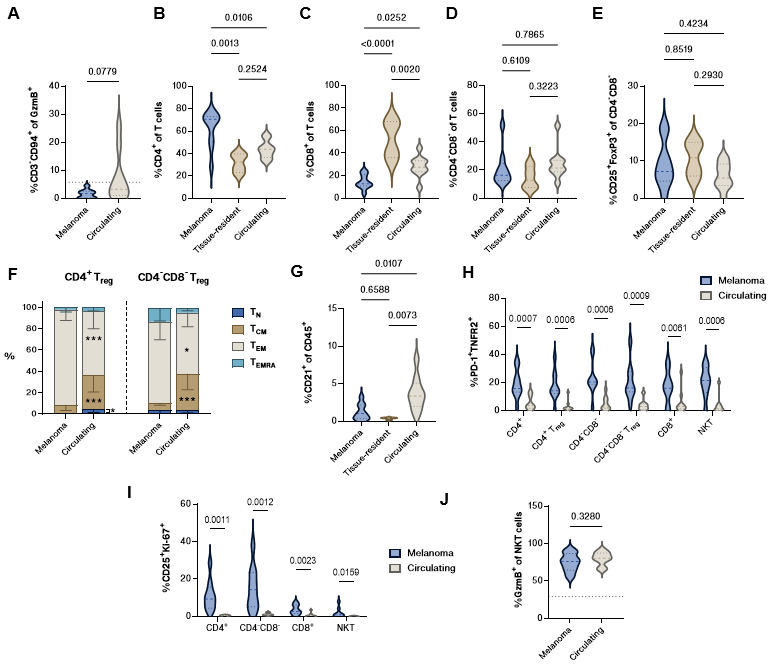

**Figure S5. Graphs comparing tumour-infiltrating, tissue-resident, or circulating immunophenotypes.** (A) Proportion of natural killer (NK) cells, defined as granzyme B (GzmB)^+^CD3^-^CD94^+^. (B-D) Proportion of CD4^+^ (B), CD8^+^ (C), and CD4^-^CD8^-^ T cells (D). (E) Proportion of CD25^+^FoxP3^+^ regulatory T cells (T_regs_). (F) Proportion of memory subsets, defined as naïve T cells (T_N_; CD62L^+^CD45RA^+^), central memory T cells (T_CM_; CD62L^+^CD45RA^-^), effector memory T cells (T_EM_; CD62L^-^CD45RA^-^), and terminally differentiated effector memory T cells (T_EMRA_; CD62L^-^CD45RA^+^), in CD4^+^ and CD4^-^CD8^-^ T_regs_. (G) Proportion of CD21^+^ B cells. (H and I) Proportion of PD-1^+^TNFR2^+^ cells (H) and CD25^+^Ki-67^+^ cells (I) within various T cell subsets. (J) Proportion of GzmB^+^CD3^+^CD94^+^ NKT cells. CD4^+^ and CD4^-^CD8^-^ represent non-T_reg_ populations. Dotted lines represent proportion in pooled oral mucosa samples. Values in bar charts represent *p*-values. Asterisks in bar charts resemble significant change with melanoma TILs; * ≤ 0.05, ** ≤ 0.01.
